# Risk Factors for Fouling Biomass: Evidence from Small Vessels in Australia

**DOI:** 10.1101/248567

**Authors:** Tracey Hollings, Stephen E. Lane, Keith R. Hayes, Felicity R. McEnnulty, Mark Green, Eugene Georgiades, Andrew P. Robinson

## Abstract

Invasive marine non-indigenous species are a major threat to marine biodiversity and marine related industries. Recreational vessels are recognised as an important vector of non-indigenous species translocation, particularly the secondary translocation of species domestically.

This paper reports on a novel application of multilevel modelling and multiple imputation to biomass samples gathered from the hull and other external surfaces of recreational yachts and fishing vessels in order to quantify the relationship between the wet biomass of biofouling and vessel-level characteristics. Unsurprisingly, we find that the number of days since the vessel was last cleaned was strongly related to the wet weight of biomass. The number of days since the vessel was last used was also related to the wet weight of biomass, yet differed depending on the vessel type. Similarly, the median number of trips undertaken by the vessel was related to the wet weight of biomass, and varied according to the type of antifouling paint used by the vessel. The relationship between vessel size, as measured by hull surface area, and wet weight biomass per sample unit area was not significant.

In order to reduce the international and domestic spread of invasive species, small vessel owners should use an appropriate type of antifouling paint that suits their vessel’s operational profile, and be encouraged to maintain a maintenance

## 1 Introduction

Invasive marine non-indigenous species are a major threat to marine biodiversity leading to dramatic shifts in population, community and ecosystem dynamics, and many have serious environmental and economic implications (Carlton and Geller, 1993; Ruiz et al., 1997; Vitousek et al., 1996). Vessel biofouling is a significant pathway for the introduction and spread of marine non-indigenous species globally (Molnar et al., 2008; Georgiades and Kluza, 2017), effectively removing natural barriers to dispersal, including ocean currents and distance (Paini and Yemshanov, 2012; Ruiz et al., 1997). Since Carlton and Geller (1993), knowledge of invasion processes and pathways for non-indigenous species in marine environments has developed considerably. International policies have been developed to regulate the ballast water pathway (International Maritime Organization, 2017) and guidelines have been developed to minimize the spread of biofouling (International Maritime Organization, 2011; International Maritime Organization, 2012). Biofouling regulations have recently been put in place by New Zealand, California and Western Australia. Because international regulation of the biofouling pathway is still relatively new, the domestic spread of non-indigenous species via this pathway, though recognised (Inglis et al., 2013; Sinner et al., 2013), remains largely unregulated. The secondary spread of non-indigenous species will ultimately determine the magnitude of economic and environmental damage caused by invasive species (Zabin et al., 2014).

The role of recreational vessels as vectors for the introduction (Ashton et al., 2014; Clarke Murray et al., 2011; Minchin et al., 2006) and spread (e.g. Burgin and Hardiman, 2011; Clarke Murray et al., 2011; Hewitt et al., 2007; Lacoursière-Roussel et al., 2012a; Minchin et al., 2006) of non-indigenous species has long been recognised. Impacts from invasive marine non-indigenous species are likely to increase with increasing use of the marine environment; for example, via the growth of marine aquaculture and increases to ship-borne trade or development of new trade routes (Hulme, 2009; Food and Agriculture Organization of the United Nations, 2016; International Maritime Organization, n.d.). In Australia, impacts from secondary invasions may become more likely as recreational vessel ownership continues to increase. In the decade from 1999–2009 vessel ownership increased by more than 36% to over 800,000 individual registered vessels nationwide (Burgin and Hardiman, 2011).

Recreational vessels have been implicated in several high profile and significantly damaging incursions of non-indigenous species including the introduction of the black striped mussel (*Mytilopsis sallei*) into a recreational marina in Darwin, Australia in 1999. Strong evidence suggests it was introduced by hull fouling on an international yacht, and detections of the mussel outside of the marina were traced to movement of smaller vessels. *M. sallei* has similar ecological and morphological characteristics to the freshwater zebra mussel (*Dreissena polymorpha*) and, at the time of discovery, densities were as high as 24,000 per square metre (Thresher, 1999). The Darwin incursion was successfully eradicated due to the fact that the mussel was contained within marinas with lock gates due to the 8 metres tidal fall (Willan et al., 2000). This situation is unlike the majority of eradication attempts of other marine non-indigenous species that have ultimately failed often despite concerted efforts. The inherent difficulties of marine invasion control has led to a focus on reducing the human-mediated spread of non-indigenous species from major ports via high risk transport vectors (Drake and Lodge, 2004; Floerl et al., 2005; Forrest and Hopkins, 2013; Georgiades and Kluza, 2017). Recreational craft may constitute a significant portion of these vectors (Inglis et al., 2012).

Biofouling is a significant pathway for introduction of marine non-indigenous species for both commercial and recreational vessels (Acosta et al., 2010; Hewitt et al., 2004; Inglis et al., 2012; Minchin et al., 2006; Thresher, 1999). Studies of international and domestic recreational vessels have demonstrated a range of fouling rates, with most surveys showing that over 50% of vessels, and sometimes as high as 80%, having some level of fouling (Ashton et al., 2014; Floerl et al., 2005; Lacoursière-Roussel et al., 2012b; Minchin et al., 2006; Zabin et al., 2014). The extent of fouling on any individual vessel, however, is often highly variable, irrespective of sample sizes and methodology (Ashton et al., 2014; Brine et al., 2013; Clarke Murray et al., 2011; Floerl et al., 2005). Some areas on a vessel are inherently more susceptible to fouling due to different hydrodynamic forces, susceptibility to coating system wear or damage, or being inadequately (or not) painted (International Maritime Organization, 2011). These “niche” areas contribute to fouling variability as they can have considerably higher levels of fouling than the hull (Lacoursière-Roussel et al., 2012b) and include internal water systems and intake pipes, rudder, propeller, and anchors/chain cabinets (Acosta et al., 2010; Burgin and Hardiman, 2011; Hewitt et al., 2007; Minchin et al., 2006).

Characteristics of recreational vessels that increase both the rate and extent of fouling are substantially different to those of commercial vessels (Lacoursière-Roussel et al., 2012b). Small vessels can have irregular maintenance and cleaning schedules, may spend significant amounts of time in port where opportunities for colonisation are high, travel at low speeds, and have varied presence and abundance of niche areas (Brine et al., 2013; Clarke Murray et al., 2011; Floerl et al., 2005; Minchin et al., 2006). Considerable variation in vessel usage and maintenance means there is equally likely to be variation in transport frequency and likelihood of non-indigenous species spread (Floerl et al., 2005). Recreational vessels often visit pristine and often protected marine areas which may be highly susceptible to invasive marine species and disconnected from commercial shipping ports and aquaculture (Clarke Murray et al., 2011; Georgiades and Kluza, 2017). Further, recreational vessels pose an additional risk of non-indigenous species transfer to aquaculture sites (Sim-Smith et al., 2016).

Vessel movements are not without risk; minimising translocation of species requires a relevant and transparent risk based profiling and assessment (Campbell and Hewitt, 2011); such assessments have been proposed in Australia as part of regulatory interventions to manage biofouling risk (Piola and McDonald, 2012; Clarke et al., 2017; Lott and Rose, 2016). If specific attributes of small vessels can be identified which increase fouling biomass and therefore facilitate the movement of non-indigenous species, then management actions and regulations can be implemented by authorities to could reduce their spread and impacts.

Using modern statistical modelling, this paper aims to identify characteristics of recreational vessels which potentially increase the extent of fouling biomass. While several studies have estimated fouling extent on a measure or index of in *situ* fouling cover (e.g. Ashton et al., 2006; Brine et al., 2013; Floerl et al., 2005), this paper offers an alternative methodology to determine vessel characteristics which influence fouling extent; that is, on directly measured fouling biomass.

Moreover, the methods which we use deal directly with the complexities of the two-stage sampling approach used to collect the data. Our data comprises samples collected from the hull and niche areas of small vessels from five locations in south-east Australia, along with vessel-level characteristics, including transport and maintenance history. The resulting data consists of two levels: multiple wet weight biomass measurements are collected for each vessel. Complicating analysis is the fact that some vessel-level characteristics are missing.

The rest of the paper proceeds as follows: in Section 2 we provide details on the data and the statistical models used for analysis; in Section 3 we present the results in detail for the full modelling procedure; and in Section 4 we provide a discussion of the results.

## 2 Methods

### 2.1 Data

The data used in this paper came from an exploratory survey in Australia conducted by the Commonwealth Scientific and Industrial Research Organisation’s (CSIRO) Centre for Research on Introduced Marine Pests (CRIMP). Biomass samples were collected from 54 vessels^1^ at 5 geographical locations during the project. All vessels were selected based on convenience when pulled out of the water for maintenance; access and permission was gathered from owners via the Bosun and slipway managers at each of the sites.

Samples were collected from both external and internal surfaces using plastic and metal putty knives, immediately following the vessels removal from the water. All samples were washed with 0.2*μ*m filtered seawater, sieved to remove as much excess water as possible, and subsequently weighed using Sartorius BL3100 scales. Due to the unreliability of low wet weight measurements, samples that weighed less than 1.5 g were assigned a biomass of 0.5 g; that is, the lower limit of detection was 1.5 g.

Most vessels were sampled in Hobart, Tasmania, Australia at the Royal Hobart Yacht Club (17) and the Domain Slipyard (16). The remaining vessels were sampled in Melbourne, Victoria, Australia at the Sandringham Yacht Club (14), the Hobsons Bay Yacht Club (4), and the Royal Yacht Club of Victoria (2). Most of the sample vessels were yachts (31), followed by commercial fishing vessels (12), motor cruisers and others (10).

Samples were collected from as many as 64 different locations in and around the hull, propeller, rudder and anchor, internal spaces, fishing gear and deck of the vessels. Samples taken from the hull^2^, fixed keel and rudder surface were consistently collected using 0.5m^2^ quadrats, so these form the focus of our analysis. There were 252 samples collected from the hull, 94 from the rudder surface, and 74 from the fixed keel; in all, 420 samples were taken. The number of samples collected per vessel ranged from 6 to as many as 10; most vessels (41) were sampled 8 times.

Vessel-level data collected from owners included: dimensions of the vessel; the type of antifouling paint applied (ablative, hard, and self-polishing); voyage history (number of days since the vessel was last used, median number of trips per year); the number of days since the vessel was last cleaned or antifouled; the surface area of the hull (m^2^); and the type of vessel (yacht, fishing, motor cruiser and others). Table 1 shows summary statistics for these observed characteristics.

For more details on the data collected during the survey, see Australian Government Natural Heritage Trust Project 46630 (Hayes et al., 2007).

**Table 1:** Summary statistics for vessel-level characteristics. Categorical variables are shown as n (%), and numeric variables are shown as median (interquartile range, IQR).

|  | Variable | n (%) / median (IQR) |
| --- | --- | --- |
| **Vessel type** |  |  |
|  | Fishing | 10 (19%) |
|  | Motor cruiser/Other | 12 (23%) |
|  | Yacht | 31 (58%) |
| **Paint type** |  |  |
|  | Ablative | 21 (40%) |
|  | Hard | 12 (23%) |
|  | Self-polishing | 15 (28%) |
|  | Missing | 5 (9%) |
| **Numeric variables** |  |  |
|  | Hull surface area (m<sup>2</sup>) | 42.0 (28.0, 57.0) |
|  | Days since last used | 12.0 (4.0, 78.0) |
|  | Days since last cleaned | 249.0 (79.0, 352.0) |
|  | Trips per year | 17.2 (10.4, 45.6) |

### 2.2 Statistical Analysis

Figure 1 shows histograms of the observed wet weight of biomass from the samples. Samples that were below the limit of detection (1.5 g, of which there were 101) are not shown in this figure. The left column shows the measured wet weight, whilst the right column shows the log-transformed wet weight; the rows correspond to the sampled locations on each vessel (hull, keel and rudder).

**Figure 1:**
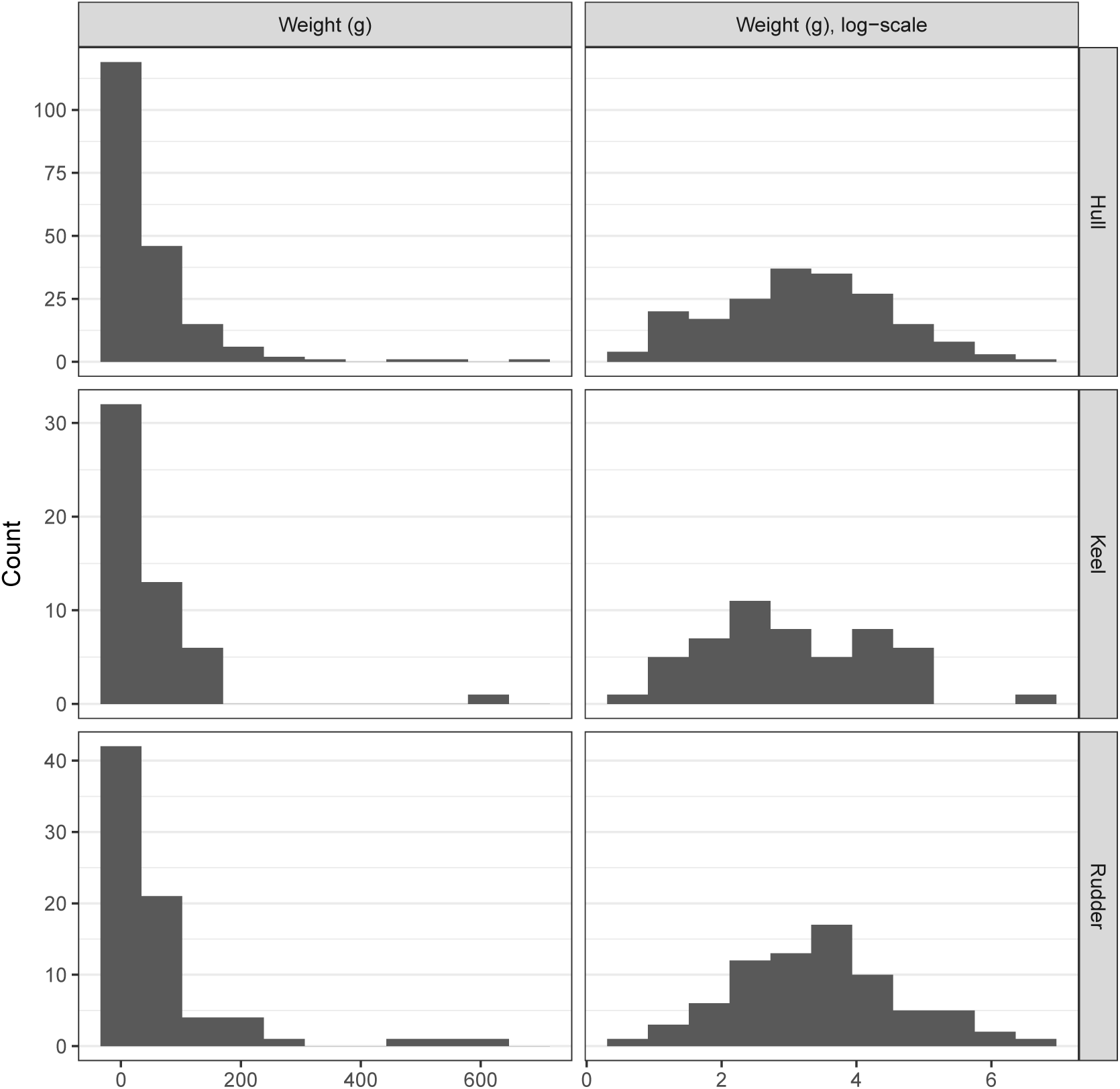
Measured wet weight of sampled biomass from the 53 vessels. The left column shows the measured wet weight, whilst the right column shows the log-transformed wet weight. Samples that measured less than the limit of detection (1.5 g) have been removed.

Figure 1 suggests that the log-transform of the wet weight of biomass has an approximately Normal distribution, and that there is likely to be a location effect (most notably between Hull versus Keel and Rudder). We thus focus on a log-transform of the wet weight of biomass for the remainder of the analysis. Further, there appears to be the possibility of outliers, especially within the hull and rudder samples. Hence we will also consider a Student-t distribution for modelling.

It could be tempting to analyse the total (or average) wet weight biomass per vessel, as this would simplify the modelling procedure. However, as noted above, Figure 1 suggests that the wet weight of biomass varies with the location from which a sample was taken; hence averaging will mask this variation. Further, the variable number of samples taken per vessel (Section 2.1) means that the total biomass may be dependent upon the number of samples taken. For these reasons, we chose to use a multilevel regression model, in which a modelled intercept allows for the repeated measurements within vessels, whilst also allowing investigation of vessel-level relationships.

To allow for the possible effects of outliers we examined the effect of changing the outcome (or observation) model from Normal to Student-*t*. Let *Y_i_* be the log wet weight of biomass for the *i*^th^ observation, *i* = 1,…, 420. Then:

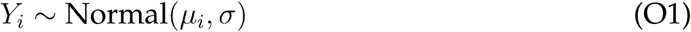

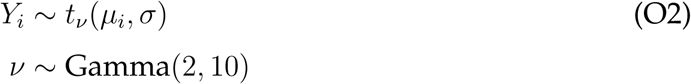
are two alternative models for the outcomes, where O1 is the model assuming a Normal distribution for log weight biomass, and O2 is the model assuming a *t* distribution with *ν* degrees of freedom. The hyperparameters for the degrees of freedom were chosen to be weakly informative, favouring a robust *t* model with lower degrees of freedom; indeed, the mode of the prior distribution for the degrees of freedom is 10.

We initially inferred the latent process — that is, the process that determines the log wet weight of fouling biomass on each vessel — as a linear function of the location of the measurement location allowing for a vessel-level intercept:

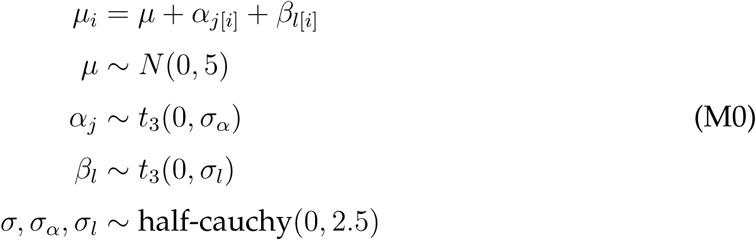
where *μ_i_* is the expected log wet weight in grams; *μ* is the intercept for the model; *β_l_* are coefficients that measure the effect of location on the log wet weight of fouling biomass: *l* = 1 denotes the measurement was taken at the hull, *l* = 2 denotes the measurement was taken at the rudder, and *l* = 3 for measurements taken at the keel^3^. *α_j_* is the vessel-level intercept that allows the hull intercept to vary from vessel to vessel, *j* = 1,…, 53, which is modelled as a *t* distribution with 3 degrees of freedom, and scale *σ_l_*. The intercept is given a weakly informative *N*(0,5) prior. Similarly the regression coefficients are given weakly informative *t*_3_(0,1) priors; as are the observation model and latent process scale parameters using half-Cauchy(0, 2.5) priors.

We subsequently investigated two alternative specifications for the vessel-level intercept by including various vessel-level characteristics and interaction terms:

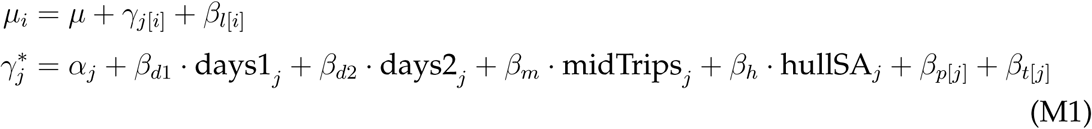

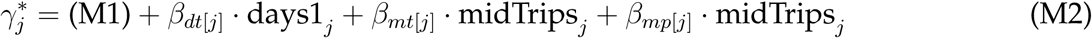

M1 includes all vessel-level characteristics, where days1*_j_* is the number of days since vessel *j* was last used; days2*_j_* is the number of days since the vessel was last cleaned; midTrips*_j_* is the median number of trips undertaken by the vessel per year; and hullSA*_j_* is the hull surface area. *β_p_*_[_*_j_*_]_ is the effect of paint type *p* on vessel *j*, *p* =1 denotes ablative paint, *p* = 2 denotes hard paint, and *p* = 3 denotes self-polishing antifouling paint. *β_t_*_[_*_j_*_]_ is an effect for vessel type; *t* =1 denotes motor cruiser/other, *t* = 2 denotes fishing vessels, and *t* = 3 denotes yachts. M2 adds interaction terms to M1 between: the number of days since the vessel was last used and the vessel type (*β_dt_*_[_*_j_*_]_); the median number of trips and the vessel type (*β_mt_*_[_*_j_*_]_); and the median number of trips and the paint type (*β_mp_*_[_*_j_*_]_).

The choice of these regression models was guided by both subject matter/process considerations and graphical exploration of the posterior distribution of the random-effect terms in the latent process models. For more details on the graphical exploration, see Section 3.

### 2.3 Missing Data

There is missing data at the vessel level; Table 2 displays the observed data pattern, shown as (Days since last used, Paint type, Days since last trip), where a 1 denotes the variable is observed, and a 0 denotes the variable is missing. For example, Table 2 shows there are 41 observations with complete vessel-level data, and 1 observation that is missing all measurements of days since last used, paint type, and days since last trip.

A complete-case data analysis would only utilise 77.4% of the vessel-level data, and subsequently, this would result in 324 (77.1%) measurements of wet weight biomass being available for the complete-case analysis at the measurement (first) level.

Because of the amount of missing data, we used multiple imputation to impute the missing vessel-level data. Specifically, we used iterative conditional imputation, with all vessel-level variables used in the imputation specification; to account for any nonlinearities, random forest imputation was used.

As all missing data was at the vessel-level, we used the median wet weight of biomass as a predictor in the imputation models (Gelman and Hill, 2007). To account for the censoring in the wet weight of biomass (observations below the 1.5 g limit of detection), we replaced the censored data by a random draw from a Uniform(0, 1.5) distribution. We justify this procedure in the following way: i) some level of variability is required as we don’t know the true value of the censored observations; ii) Figure 1 shows that weights below 1.5 g (0.4 on the log-scale) form a very small part of the range of observed values. This small variability will have limited impact in the imputation of the missing data due to the fact that we are summarising within-vessel wet weight of biomass measurements by the median for input into the imputation models.

To perform inference, we follow the advice of Zhou and Reiter (2010) and mix the draws from the posterior distributions from the analysis of each completed dataset^4^. Zhou and Reiter (2010) recommend a large number of imputations be used; we generated 50 imputed datasets. For each dataset, 2000 samples were drawn from the posterior after a burn-in of 2000. Convergence was monitored by inspecting the mixed chains across the 50 multiple imputations, and by assessing the Gelman-Rubin statistic (Gelman and Rubin, 1992).

**Table 2:** Observed data pattern for vessel-level characteristics. The patterns are shown as (Days since last used, Paint type, Days since last trip), where a 1 denotes the variable is observed, and a 0 denotes the variable is missing. A count of each type of pattern is shown.

| Observed data pattern | Count |
| --- | --- |
| (1, 1, 1) | 41 |
| (0, 1, 1) | 1 |
| (1, 1, 0) | 5 |
| (1, 0, 1) | 4 |
| (0, 1, 0) | 1 |
| (0, 0, 0) | 1 |

<sup>4</sup>The alternative would be to combine posterior estimates of summary parameters using Rubin's rules (Rubin, 1996), which necessarily gives point estimates as opposed to full posterior distributions.

All analyses were performed using R version 3.4.0 (R Core Team, 2017); the multilevel models were fit using the RStan interface to Stan (Stan Development Team, 2016), and multiple imputation was performed using the R package mice (Buuren and Groothuis-Oudshoorn, 2011). R code and data to reproduce the analysis may be found at https://github.com/SteveLane/blistering-barnacles.

## 3 Results

### 3.1 Multiple Imputation

Visual comparison between the observed and imputed data suggests that the imputation procedure has satisfactorily converged, i.e. there was no large discrepancies between the imputed data and the observed. The comparison between observed and imputed data is presented in the Supplemental Material.

### 3.2 Outcome Model

The *t*-distribution observation model (O2) outperformed the Normal distribution observation model (O1). The difference in the leave-one-out information criterion (LOOIC, analogous to AIC, see Vehtari et al., 2016) was 47 (standard error 13) in favour of the *t*-distribution observation model (O2).

Posterior predictive checks likewise confirmed the superiority of the *t*-distribution observation model. Graphical checks were made using the proportion of posterior predictions that lie below the limit of detection; the median posterior predictions and the interquartile range (IQR) of the posterior predictions. The graphical checks and LOOIC comparisons can be found in full in the Supplemental Material.

### 3.3 Regression Results

#### 3.3.1 Exploration of Candidate Interaction Terms

Figure 2 displays the posterior median and 80% credible intervals of the regression coefficients in the vessel-level intercept models (for each of the four ordinal predictors — the number of days since the vessel was last used; the number of days since the vessel was last cleaned; the median number of trips undertaken by the vessel; and the vessel’s hull surface area). Figure 2a shows the coefficients coloured by vessel type, whilst Figure 2b shows the coefficients coloured by antifouling paint type. Each panel also shows the posterior (median) estimate of the expected log wet weight from model M1, and is based on a randomly selected imputed dataset for display purposes.

**Figure 2:**
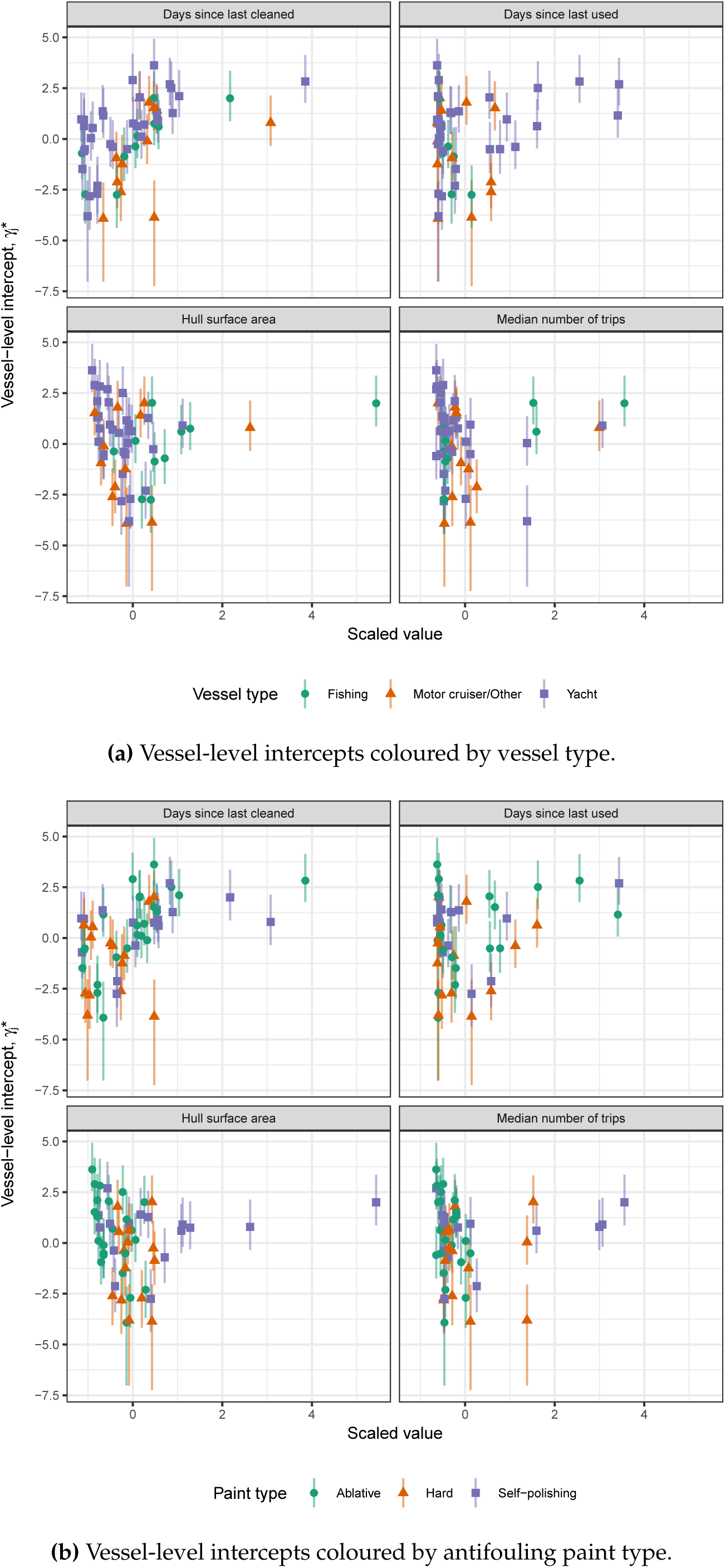
Estimated vessel-level intercepts from M1, plotted against the four ordinal predictors. Each plot shows the median and 80% credible interval, coloured by vessel type (2a) and paint type (2b). The (median) regression lines for each predictor are overlaid.

Figures 2a and 2b are suggestive of interactions between the number of days since the vessel was last used and the vessel type, the median number of trips undertaken by the vessel and the vessel type, and the median number of trips taken by the vessel and the antifouling paint type. We form this opinion by visual comparison of the scatterplots. For example, the pattern in Figure 2a suggests that the effect of days since the vessel was last used is negative for fishing vessels and positive for yachts and motor cruiser/other vessels. Similarly, the effect of the median number of trips undertaken by the vessel appears to be positive for fishing vessels, and negative for yachts and motor cruiser/other vessels.

#### 3.3.2 Final Model Form

Figure 3 displays the median and 80% intervals of the estimated vessel-level intercepts, with respect to the two ordinal predictors used in interaction terms in (M2): the number of days since the vessel was last used and the median number of trips (per year) undertaken by the vessel. Figure 3a shows the vessel-level intercepts coloured by vessel type and Figure 3b shows the vessel-level intercepts coloured by antifouling paint type. Shown in each panel are the estimated (median) regression lines from M2.

**Figure 3:**
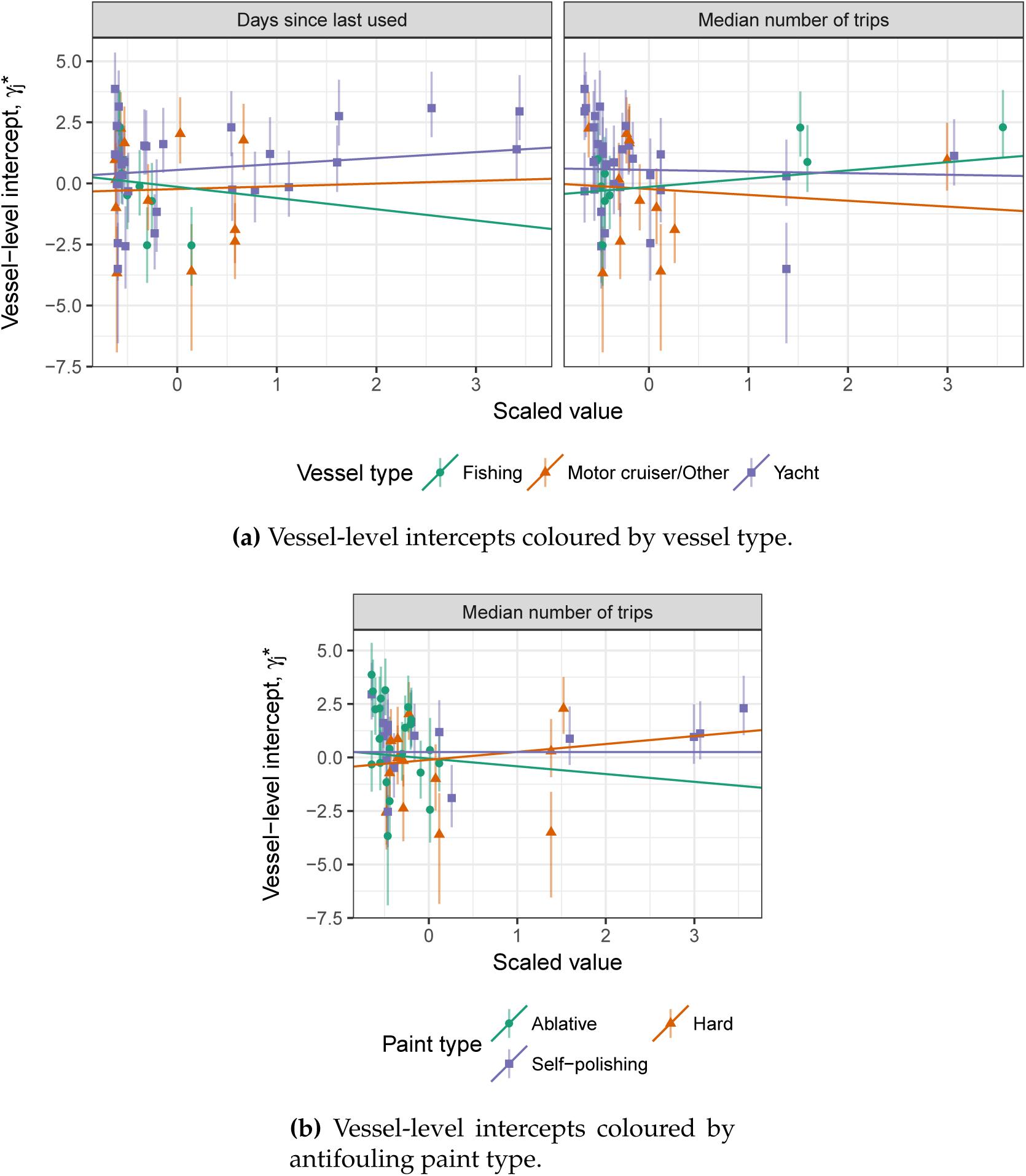
Estimated vessel-level intercepts from M2, plotted against the two ordinal predictors with interactions being assessed. Regression lines for each estimated interaction are overlaid.

Figure 3a suggests that the of the number of days since the vessel was last used varies by vessel type, as does the median number of trips taken by the vessel per year. Figure 3b further suggests that the effect of the median number of trips taken by the vessel per year varies according to the paint type of the vessel. The uncertainty in these effects is however, large relative to the estimates.

Figure 4 compares estimates of the measurement-level and vessel-level regression coefficients for each of the models considered. As can be seen in Figure 4, the effect of the number of days since the vessel was last cleaned (*β_d_*_2_) is clearly larger than 0; the other predictors do not appear to have such a strong effect. Whilst the effect of these predictors is not estimated precisely, the direction and size of the coefficients is similar between models, and as such we choose Model (M2) as the final model form in order to estimate the interactions.

**Figure 4:**
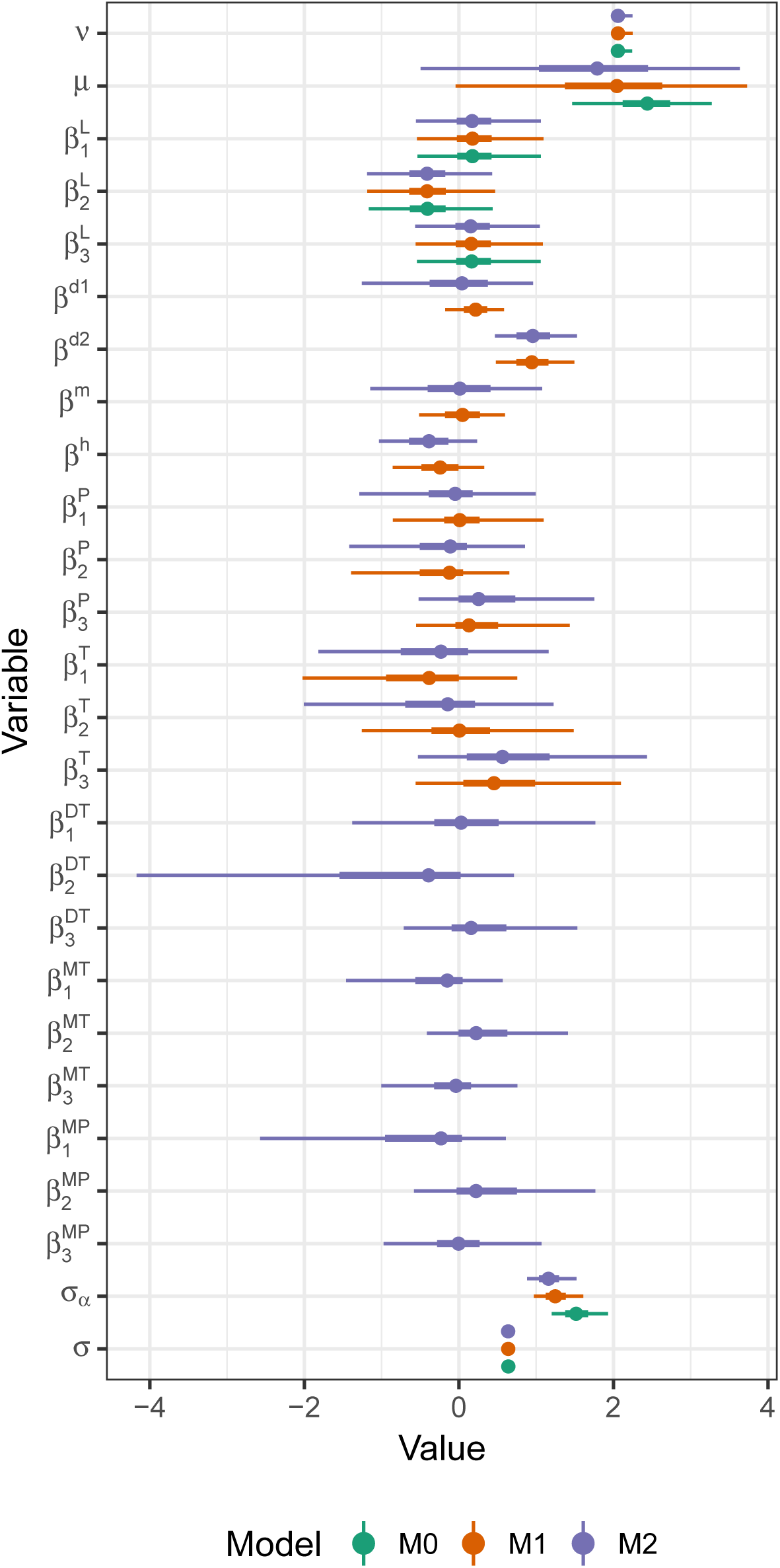
Estimated coefficients from M0–M2. Coefficients from each model (where applicable) are shown grouped within coefficient on the y-axis. The median is shown as a point estimate, with 50% and 90% credible intervals.

#### 3.3.3 Predictive Model Comparison

Table 3 compares the models by their leave-one-out information criterion (LOOIC, analogous to AIC, see Vehtari et al., 2016). M2 is the best performing model based on LOOIC, but the variability is sufficiently large that all models are closely performing. The difference in LOOIC between M0 and M2 (3.5) is almost one and a half times the standard error of the difference.

**Table 3:** LOOIC-based comparison of models M0–M2. The model with the smallest LOOIC (M2) is shown first, with subsequent rows ordered by increasing LOOIC (smaller LOOIC values are preferred). ΔLOOIC shows the difference in LOOIC between all models and M2; se(ΔLOOIC) shows the estimated standard error of the difference. Eff. P gives the estimated effective number of parameters; se(Eff. P) shows its standard error.

| Model | LOOIC | $se(\text{ELPD})$ | Eff. P | $se(\text{Eff. P})$ | ΔLOOIC | $se(\Delta$LOOIC) |
| --- | --- | --- | --- | --- | --- | --- |
| M2 | 1102 | 21.7 | 70.2 | 4.29 |  |  |
| M1 | 1103 | 21.6 | 70.5 | 4.29 | 0.36 | 0.827 |
| M0 | 1106 | 21.7 | 73.3 | 4.90 | 3.55 | 2.522 |

## 4 Discussion

Several previous studies and vessel profiling by regulatory authorities have proposed vessel characteristics that should be indicative of biofouling translocation risks, including voyage time, time in the water, time since application of antifouling paint, frequency of vessel use and stationary period (Ashton et al., 2006; Hewitt et al., 2011; Ministry for Primary Industries, New Zealand, 2016; California State Lands Commission, 2017; Department of Fisheries, Western Australia, n.d.).

Some analyses, however, have either failed to find a strong relationship between vessel characteristics and fouling extent or abundance, or provided contradictory evidence. Clarke Murray et al. (2011) for example, found percent fouling cover on recreational yachts was not related to travel frequency or the age of antifouling paint. Floerl et al. (2005) concluded that the abundance of non-indigenous species on yacht hulls could not be associated with maintenance schedules, travel history or owners maintenance behaviour. Similarly, Ashton et al. (2014) did not find significant correlations between last cleaning and extent of fouling using in-water surveys, and also concluded that the type of recreational vessel does not explain differences in biofouling patterns.

Other studies however, suggest that time since the antifouling paint was applied to the vessel is a consistently reliable predictor of biofouling (Ashton et al., 2006; Floerl et al., 2005; Lacoursière-Roussel et al., 2012b; Ashton et al., 2014), and that time in the water, stationary periods, and voyage types are relevant factors (Ashton et al., 2006; Lacoursière-Roussel et al., 2012b). Lacoursière-Roussel et al., 2012b also suggest that vessel type does play a role because powerboats and catamarans, for example, have a greater risk of fouling due to their higher number of niche areas.

The results presented here demonstrate relationships that are largely consistent with prior expectations about important risk factors; that is, the amount of hull fouling biomass is related to the time since a vessel was last cleaned and last used, how frequently a vessel is used and the type of antifouling paint applied. Consider for example a yacht that hasn’t been cleaned in 249 days. The model predicts that for such a vessel, there is a 74% probability that delaying cleaning by six months would result in an increase of wet weight biomass within a 0.5m^2^ area on the hull of the vessel. This result is in agreement with recommendations of continual maintenance by the Australian and New Zealand Antifouling and In-water Cleaning Guidelines (Department of Agriculture, Canberra, 2015; Georgiades et al., 2018).

The size of the vessel, measured by the hull surface area, was found to have a small negative relationship if used to predict biofouling levels; however, the uncertainty in this estimate was relatively large. Given the same frequency of use, the modelling suggests that different types of vessels have different effects on biofouling levels. For example, assume that vessels have the same duration since last being cleaned, and the same duration since last being used; the model predicts that there is an 72% predicted probability that a yacht will have a larger wet weight biomass within a 0.5m^2^ area than the motor cruiser/other vessel; compared to a fishing vessel, this probability is 75%.

A weak varying effect of the median number of trips taken by the vessel per year and paint type was found in the model. The limited number of data points at the vessel level (53) hampered the precision of estimating these effects, but the direction of effects conforms with prior expectations, namely that: both ablative and self-polishing paints are reliant on the vessel being used in order to be effective, whilst hard paints are less so (Georgiades et al., 2018).

There is also evidence that the location of the sample is an important predictor for the wet weight of biomass. Figure 4 shows a consistent result that the wet weight of biomass when sampled along the hull 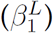, is no different to samples made from the rudder 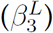. Measurements made on samples from the keel however, are generally smaller than those made along either the hull or on the rudder; in fact, there is a 69% predicted probability of this being the case. This is not a characteristic of vessels, and thus cannot be used for management purposes; the location of the measurement does however need to be considered in any future work, and in particular may restrict vessel-level summaries of wet weight of biomass being analysed (for example, total wet weight of biomass).

Fouling cover on vessels can be up to ten times higher in niche areas than on hulls (Clarke Murray et al., 2011), and these areas represent important settlement locations of non-indigenous species (Lacoursière-Roussel et al., 2012b; Zabin et al., 2014). Floerl et al. (2010) and Ashton et al. (2014) agree on the importance of targeting niche areas for vessel surveys and sampling.

This study is not without its limitations. We found that the (log-transformed) wet weight of biomass found on small vessels was best described by a t-distribution. Posterior predictive checks showed that the analysis regression models generated data consistent with the observed data for the interquartile range, but less so for the median and proportion of observations below the limit of detection. A possible approach to allow for the larger number of censored observations than expected would be to fit a mixture model. It is not directly clear how this would work however, when vessels may contain some samples measured below and some above the limit of detection. Further, given the limited data available, it was considered that an attempt to fit such a mixture model was unfeasible.

The small number of vessels sampled also limits the precision at which we can estimate the effect of vessel-level characteristics, and in particular, any interactions between them. We demonstrated a number of possible interaction effects, yet the error in those estimates was found to be relatively large. The nature of these varying effects however, did conform to prior expectations of the direction of the effect; it is the size of such effects that is not known with a high degree of certainty.

Sample size reservations notwithstanding, we have confirmed that vessel-level characteristics can be used to predict the amount of wet weight of fouling biomass that may be present on small vessels in Australia. The development of such predictive tools could aid authorities in assessing the risk posed by individual vessels and classifying those that are a risk of having high levels of fouling will potentially reduce the introduction and spread of invasive species (Minchin et al., 2006; Clarke et al., 2017).

The results suggest that in particular, owners of small vessels should be encouraged to apply an antifouling paint that is appropriate to their vessel operational profile and to maintain a regular cleaning and maintenance schedule in accordance with the manufacturer’s instructions as per Australian and New Zealand guidance (Department of Agriculture, Canberra, 2015; Georgiades et al., 2018). Management interventions, such as diver inspections, directive to clean at a dry dock etc, can be implemented to varying degrees based on increasing levels of risk, but also with a prohibitive increase in resources, cost and time (Campbell and Hewitt, 2011). Therefore strategies and incentives could be investigated to encourage vessel owners to improve their cleaning and maintenance, as well as education and outreach programs for vessel owners to increase awareness of invasive species and their impacts (Brine et al., 2013; Minchin et al., 2006).

Authorities could also introduce regulated antifouling paint regimes, e.g. every 12 months, regular inspections and providing cleaning facilities for recreational vessel users (Zabin et al., 2014).

Preventing the introduction of invasive marine species is the key to reducing their spread, requiring stringent implementation of measures to reduce invasions through all vector routes (Bax et al., 2001; Inglis et al., 2013; Sinner et al., 2013). While interregional movements of vessels are not without risk, minimising species translocations will require a relevant and transparent risk based profilinf and assessment (Campbell and Hewitt, 2011; Clarke et al., 2017), taking into account the characteristics of transport vectors (Floerl and Inglis, 2005). Such profiling and assessment would be assisted by promoting ongoing maintenance and consistent “clean before you leave” messaging within a broader domestic pathway management approach.

## Acknowledgements

The authors would like to acknowledge Dr Enamul Kabir for his contribution to earlier analyses that were not a part of this manuscript, and to Sandringham College, Melbourne, Victoria, for use of the school’s laboratory space to process the samples taken within Melbourne. AR, SL and TH’s contributions were funded in part by the Centre of Excellence for Biosecurity Risk Analysis.

## Footnotes

1 One vessel was removed from analysis due to its abnormally high biomass measurements.

2 Fore, midships and aft were all sampled, both port and starboard.

3 The notation *l*[*i*] denotes the location level (*l*) of the *i*th observation. The vessel-level intercept (*α_j_*_[_*_i_*_]_) is similarly referenced.

4 The alternative would be to combine posterior estimates of summary parameters using Rubin’s rules (Rubin, 1996), which necessarily gives point estimates as opposed to full posterior distributions.

